# Influence of plant genotype on nodule microbiome composition and function in seasonal yellow pea varieties

**DOI:** 10.1101/2025.09.24.678337

**Authors:** Henrique M. Dias, Shyam Solanki, Jose L. Gonzalez Hernandez, Ravi V. Mural, Svetlana N. Yurgel, Christopher Graham

## Abstract

**Background and Aims:** Legume root nodules host symbiotic rhizobia essential for nitrogen fixation but also harbor diverse non-rhizobial taxa that remain poorly characterized. Yellow pea (*Pisum sativum*) cultivars adapted to distinct seasonal growth (spring and winter) offer an opportunity to explore whether host genotype influences nodule microbiome composition and function. This study investigates the taxonomic and functional profiles of nodule-associated microbial communities in seasonal yellow pea varieties.

**Methods:** A field experiment with 6 field pea cultivars (spring and winter types) was conducted in South Dakota. Surface-sterilized root nodules were subjected to full-length 16S rRNA gene sequencing using Oxford Nanopore technology. Reads were quality filtered, organellar sequences removed, and taxonomic classification performed with the EMU pipeline. Microbial diversity, community structure, and core taxa were analyzed using R, with predicted functions inferred by FAPROTAX.

**Results:** The nodule microbiome was dominated by *Rhizobium*, accounting for up to 98% of classified reads. After excluding *Rhizobium*, non-rhizobial diversity revealed a conserved core microbiome shared across cultivars, including cyanobacteria with potential phototrophic and diazotrophic traits. Minor seasonal differences were observed, with winter cultivars exhibiting higher evenness and specific associations.

**Conclusion:** Yellow pea nodules harbor a stable, cyanobacteria-enriched core microbiome, largely consistent across seasonal cultivars. Season-specific microbial patterns suggest potential host-genotype influences, warranting further validation.

## INTRODUCTION

Plants have evolved close associations with soil microbes that contribute to nutrient uptake, enhance resistance to pathogens and abiotic stresses, and support expansion into new environments (Bever et al., 2012). Recent studies highlight the role of the plant microbiome in enhancing plant adaptability to changing environments by modulating phenotypic plasticity (Camargo et al., 2023). In this context, roots are the main point of contact between plants and soil, and in the rhizosphere, they support a diverse and dynamic microbial community. These communities are shaped by constant interactions between plants and microbes, as well as among microbes themselves. Many of these interactions are mediated by molecular signals, such as secondary metabolites, which can influence microbial composition, support beneficial microbes, or suppress harmful ones (Hacquard et al., 2024). Understanding what drives the assembly of these microbial communities and how plants and microbes adapt together can help guide efforts to improve crops and promote more sustainable agricultural practices (Zhang et al., 2017).

As an example of this, legume root nodules have traditionally been studied as specialized organs for the fixation of symbiotic nitrogen, inhabited primarily by rhizobia such as *Bradyrhizobium*, *Rhizobium*, and *Mesorhizobium*. These symbiotic relationships have been shown to contribute significantly to reducing the dependence on synthetic fertilizers (Zhu et al., 2023). In addition, high-resolution microbiome profiling has revealed that nodules contain more than rhizobia; they are microbiome-rich environments colonized by various non-rhizobial taxa (Han et al., 2020; Hossain et al., 2023). Genera such as *Pseudomonas*, *Enterobacter*, and *Bacillus* have been identified in nodules, with roles that extend beyond nitrogen fixation, including the promotion of plant growth, suppression of disease, and improved resilience under environmental stress (Martinez-Hidalgo et al., 2017; Yin et al., 2021; Carter et al., 2023).

Furthermore, the microbiome associated with the nodule is highly dynamic and strongly affected by the genotype of the host plant (Burghardt et al., 2022). Even under uniform environmental conditions, different legume cultivars recruit different microbial communities. For instance, *Glycine max* cultivars exhibit variation in the abundance of non-rhizobial taxa such as Pseudomonadaceae and Enterobacteriaceae (Yang et al., 2023). This characteristic suggests that host plants act as selective filters, possibly through root exudates (Zhalnina et al., 2018), immune responses (Teixeira et al., 2021) and signaling pathways that shape microbial compatibility (Reid et al., 2018). Moreover, some recent studies suggest that host genotypes not only influence microbial recruitment (Ishida et al., 2022; Dilla et al., 2021), but also affect community function, with consequences for nitrogen fixation efficiency, community stability, and stress responses (Wendlandt et al., 2019; Crosbie et al., 2022).

Like other legumes, field pea (*Pisum sativum*) can form root nodules that fix atmospheric nitrogen through interactions with host-specific rhizobia in cultivated soils (Sujkowska et al., 2011). This crop is particularly valued for its protein-rich seeds and has attracted increasing interest due to its dual utility as both a cover crop and a source of high-quality protein for food applications (USDA, 2010). Historically, winter-sown and spring-sown cultivars were selected based on their seasonal adaptability and agronomic functions. Winter peas offer early soil cover and nitrogen enrichment, while spring peas have been favored for seed production and food quality traits (USDA, 2022). These groups also exhibit contrasting physiological traits, including cold tolerance, growth duration, and flowering time, which can influence root development and rhizosphere interactions (Vocanson et al., 2008; Vann et al., 2018). Such differences raise compelling questions about whether divergent domestication pathways have shaped distinct rhizobial recruitment strategies or microbiome assemblies between field pea cultivars (Yurgel et al., 2025).

In this study, we focus on yellow field pea, a crop of agronomic importance with cultivars adapted to different seasonal growth conditions (spring and winter). Given the increasing recognition that non-rhizobial microbes can shape legume performance, we hypothesize that seasonal yellow pea varieties recruit distinct nodule microbiomes that contribute to functional variation in symbiosis. Using 16S rRNA metabarcoding, we characterized the taxonomic and predicted functional profiles of nodule-associated microbial communities in six varieties of yellow pea. We identified both core and variable microbial taxa associated with each seasonal group, offering new insights into host-microbiome interactions in legumes.

## MATERIALS AND METHODS

### Field and experimental design

The field experiment was conducted at the West River Research Farm of South Dakota State University in Sturgis, South Dakota, located at 44° 25’ 28.53" N, 103° 22’ 22.41" W. The site is characterized by a semi-arid climate (DBsk) with an average annual precipitation of 496.82 mm and a temperature of 8.4°C (Daly et al., 2013). The soil at the experimental site is classified as Nunn clay loam (NcA), with a pH in the top 30 cm of 6.4 and organic matter content of 27 g kg-1. This trial was established concurrently with annual field pea variety trials. Field peas were planted on October 3, 2022 for winter cultivars and April 27, 2023 for spring cultivars at 25-cm row spacing with a no-till grain drill (Model 750, John Deere Co., Moline, IL, USA) at a population of 86 pure live seedm−2. A commercial granular inoculant (Exceed, Visjon Biologics, Henrietta, TX, USA) was applied with the seed at manufacturers recommended rates. No additional fertility was applied. For weed control, a premix herbicide of carfentrazone ethyl and sulfentrazone was applied as a postemerge application at a rate of approximately 365 ml ha-1. The experiment was carried out on plots that have not previously had field peas grown on them using a completely randomized block design with three replications, featuring two seasonal groups of yellow pea varieties: spring and winter, consisting of three cultivars per group. Individual plots measured 1.5 m. x 9.1 m. accommodating 6 rows per plot.

### Sampling, surface sterilization and DNA extraction

At anthesis, root nodules were carefully collected from the center two rows of each plot from the crown region of five individual plants, resulting in five distinct clusters of nodules. To remove adhering soil particles, the nodules were gently washed with sterile water. Following the sterilization protocol from Sharaf et al. (2019), surface sterilization was performed by rinsing the nodules three times with a 0.9% NaCl solution, followed by rolling them on lint-free wipes to remove any remaining soil. The nodules were then soaked twice for 5 minutes in a 1.65% (v/v) sodium hypochlorite solution, with gentle inversion during each soak, and rinsed with sterile water between treatments. Finally, the nodules were rinsed three additional times with sterile water to ensure complete removal of the sterilizing agent. After sterilization, the nodules were stored at −80°C until DNA extraction. DNA was extracted from pooled nodule samples for each cultivar using the DNeasy Plant Pro Kit (Qiagen) according to the manufacturer’s instructions.

### 16S rRNA full-length amplification, library, and sequencing

The composition of the microbial community was assessed by amplifying the full-length 16S rRNA gene from 10 ng of DNA using the Bact26F forward primer (5’- AGAGTTTGATCMTGGCTCAG-3’) and the Univ1492R reverse primer (5’- GGTTACCTTGTTACGACTT-3’) (Galkiewicz and Kellogg, 2008). PCR reactions were performed in a total volume of 40 µL, containing 25 µL of LongAmp Hot Start Taq 2X Master Mix (NEB, M0533), 10 µL of nuclease-free water, and 5 µL of template DNA (25 ng). The thermal cycling conditions were as follows: initial denaturation at 95°C for 1 min; 25 cycles of denaturation at 95°C for 20 sec, annealing at 55°C for 30 sec, extension at 65°C for 2 min; and final extension at 65°C for 5 min. Following amplification, the PCR products were purified using AMPure XP beads (Beckman Coulter, A63881) according to the manufacturer’s instructions. Library preparation, including indexing, quantification, normalization, pooling, and loading, was performed following the 16S Rapid Sequencing Barcoding Protocol (Oxford Nanopore Technologies, SQK-16S024). Sequencing was performed on the PromethION platform (Oxford Nanopore Technologies) at the South Dakota State University (SDSU) Genomics Core Sequencing Facility.

### Amplicon sequence analysis

Raw sequencing reads in FASTQ format were first demultiplexed using the DORADO Fastq Barcoding workflow v3.10.4 (Oxford Nanopore Technologies). The quality assessment of the raw reads was performed with NanoStat v1.6.0, followed by quality filtering using NanoFilt v2.8.0. Reads with a Phred quality score below 12 and sequences shorter than 1,000 bp and longer than 1,800 bp were discarded to ensure high-confidence data for downstream analysis. The reads were then screened for chloroplast and mitochondrial sequences using Kraken2 v2.1.3 (Wood et al., 2019) with 0.6 of confidence using a custom Viridiplantae organelle database (RefSeq, May 5, 2025). Organellar reads were excluded prior to taxonomic classification. Subsequently, reads were processed through the EMU pipeline (Curry et al., 2022) using default parameters, for taxonomic classification and estimation of relative microbial abundances. The taxonomic assignment was performed using an EMU pre-built database that integrates NCBI 16S RefSeq database (February 1, 2023). Taxonomic annotations were retained at the species, genus, and family levels to capture a broad, yet detailed, overview of the composition of the microbial community. Only taxa with a relative abundance ≥0.001 and that were detected in at least two independent replicates were retained for all downstream analysis. The functional profiles of bacteria were predicted using FAPROTAX v1.2.11 (Louca et al., 2016) according to taxonomic assignments at the species level. The predicted functions were grouped as core, spring-specific, and winter-specific for comparative analyses.

### Data analysis

All bioinformatic and statistical analyses were performed in R (v4.1; R Core Team, 2021). The alpha diversity indices, including Shannon diversity, Simpson diversity, the Chao1 richness estimator, and Pielou’s evenness, were calculated using the *vegan* package (v2.5-7) (Oksanen et al., 2013). Beta diversity was assessed using Bray-Curtis dissimilarity, followed by principal coordinate analysis (PCoA) and non-metric multidimensional scaling (NMDS), with ordination plots visualized using 95% confidence ellipses. Differences in community composition were tested using PERMANOVA (Adonis2 function, vegan package) with 999 permutations. Taxonomic composition was analyzed at the species and family levels and grouped by seasonal cultivar (spring and winter). The core microbiome was defined as taxa present in at least 90% of cultivars within each seasonal group, with a minimum of 500 reads and detection in at least two of three replicates. To minimize bias, only taxa present in all biological replicates of each cultivar were considered for the classification of the core microbiome. Venn diagrams and Euler plots were generated to compare taxa unique to winter and spring cultivars and those shared between groups. Pairwise differences in *Rhizobium* relative abundance were assessed using Wilcoxon rank-sum tests. Differential abundance analyses at the genus level were performed using DESeq2 (v1.34.0) (Love et al., 2014), with the Benjamini-Hochberg false discovery rate (FDR) correction applied to control for multiple tests (p < 0.05 threshold).

## RESULTS

### Sequencing output and taxonomic classification

A total of 11,453,755 full-length 16S rRNA reads were generated averaging 673,750 reads per sample, with a median read length (N50) of 1,549 base pairs (bp). After basecalling, reads were filtered for a minimum quality score of 12 and lengths between 1,000 and 1,800 bp, retaining 10,714,816 high-quality reads. Host-derived organellar contamination represented 0.1% of reads and was removed prior to downstream analysis. EMU-based classification then resolved dominant taxa to the species-level, with unmapped and unclassified reads comprising no more than 0.0005% of the dataset (**Supplementary Table S1**). In total, Emu identified 197 unique species in 111 genera, 65 families, and six phyla (**Supplementary Table S2**).

### Rhizobium dominance in yellow pea nodules

To compare the composition of the microbial community between spring and winter yellow pea cultivars, we analyzed the relative abundance profiles at different taxonomic levels. At the phylum level, the nodule microbiome was dominated by Proteobacteria (approximately 96% of classified reads), followed by Cyanobacteria (approximately 3%). Other phyla, including Actinobacteria, Bacteroidetes, Firmicutes, and Synergistetes, together contributed less than 1% (**Figure 1A**). Having established the broad taxonomic landscape, we next assessed seasonal differences at the genus level. As expected from the nodular origin of the samples, *Rhizobium* was the most abundant genus, with a relative abundance ranging from 0.84 to 0.98 (**Figure 1B-C**). Although its overall dominance was consistent, hierarchical clustering based on *Rhizobium* abundance revealed intra-seasonal variability across cultivars (**Figure 1C**). For instance, while some winter cultivars such as Keystone and Vail clustered at the high end of the abundance spectrum (>0.96), spring cultivars showed comparatively variable levels. Given the incomplete season-based segregation observed in clustering, *Rhizobium* abundance appears to reflect a combination of cultivar-specific and other ecological/structural factors rather than season alone.

**Figure 1.**
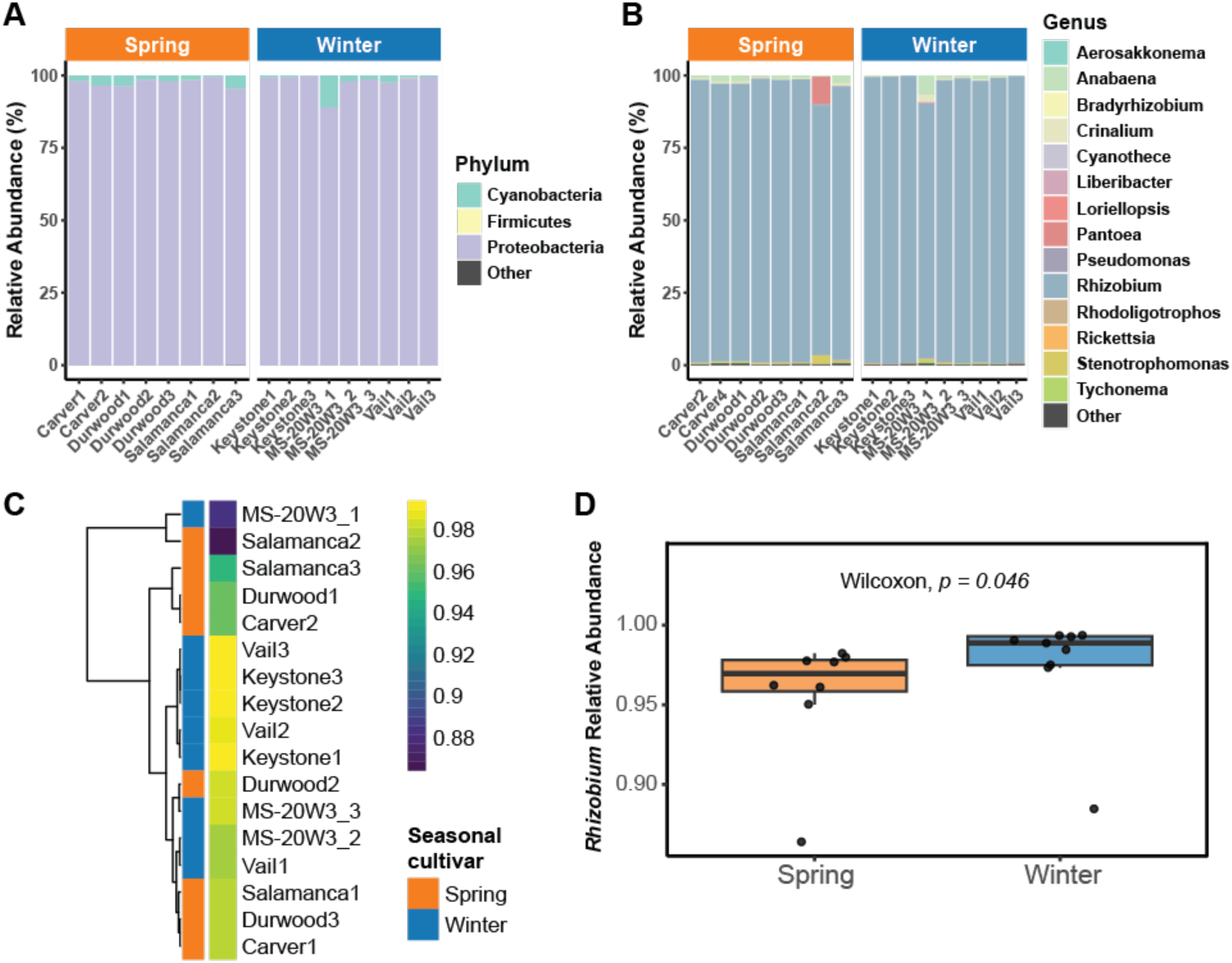
Seasonal differences in the microbial community structure and *Rhizobium* dominance in yellow pea nodules. **A)** Phylum-level composition of nodule-associated bacteria across seasonal cultivars. Stacked bar plots show the relative abundance (%) of bacterial phyla detected in individual nodule samples from spring (orange) and winter (blue) cultivars. Proteobacteria dominated the microbiome in all samples, while Cyanobacteria and Firmicutes were present at lower abundances. Sample names are labeled along the x-axis by cultivar and replicate. Bars are grouped by seasonal cultivar. **B)** Genus-level taxonomic composition of nodule-associated bacteria across seasonal cultivars. Relative abundance (%) of bacterial genera detected in each nodule sample, grouped by seasonal cultivar. *Rhizobium* is the dominant genus in both groups. Genera are color-coded as indicated in the legend. Each bar represents one replicate. **C)** Heatmap of genus-level relative abundances across samples. A heatmap displays the relative abundance of bacterial genera across spring and winter nodule samples. Rows correspond to individual samples (n = 18), and columns represent bacterial genera. The color gradient (purple to yellow) indicates an abundance of values. A dendrogram based on hierarchical clustering of genus profiles reveals patterns of similarity among samples. Colored side bars denote seasonal cultivars (orange = spring, blue = winter). **D)** Comparison of *Rhizobium* abundance among seasonal cultivars. Boxplots depict the relative abundance of *Rhizobium* in spring (n = 8) and winter (n = 9) samples. *Rhizobium* was significantly more abundant in winter nodules (Wilcoxon rank-sum test, *p* = 0.046). Boxes represent the interquartile range (IQR), and individual data points are plotted.

Finally, we formally tested whether the small seasonal difference in *Rhizobium* abundance was significant. A Wilcoxon test showed that winter cultivars harbored a modest but statistically higher median relative abundance of *Rhizobium* compared to spring cultivars (**Figure 1D**, *p* = 0.046), consistent with the clustering result yet underscoring that season only explains a small fraction of the overall variation. These results confirm *Rhizobium* as the dominant nodule taxon, with minor but potentially meaningful differences between individual cultivars that could reflect genotype-driven modulation of symbiotic recruitment, but this requires larger sample sizes for confirmation.

### Microbiome comparison between spring and winter cultivars

To evaluate potential seasonal effects on microbial diversity, we first compared alpha diversity between spring and winter yellow pea cultivars using Shannon and Simpson diversity indices, Pielou’s evenness, and Chao1 species richness. While Chao1 richness did not differ significantly between seasonal cultivars, Shannon diversity was marginally higher in spring (*p* = 0.0543), and both Simpson’s diversity and Pielou’s evenness were significantly greater in spring compared to winter (*p* = 0.0433; **Figure 2A**). Next, we assessed beta diversity using Bray-Curtis dissimilarity, visualized through principal coordinate analysis (PCoA) and non-metric multidimensional scaling (NMDS). The spring and winter cultivar samples did not form distinct groups in either ordination method, and PERMANOVA analysis confirmed no significant difference in community composition between groups (*p* > 0.05) (**Figure 2B-C**).

**Figure 2.**
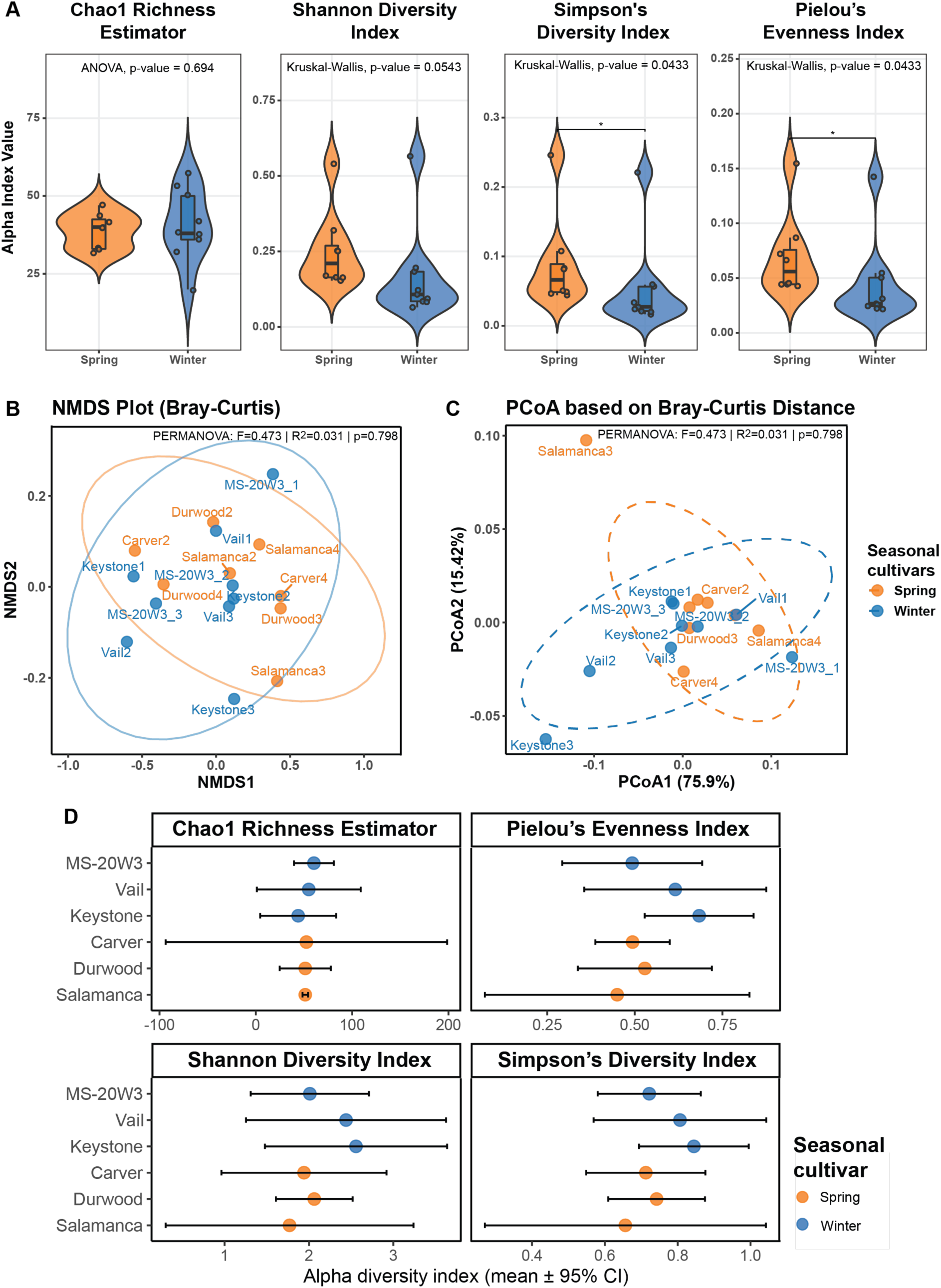
Seasonal cultivar effects on alpha and beta diversity of nodule-associated bacterial communities. **A)** Alpha diversity indices of spring and winter cultivars at genus-level. Violin plots show distributions of four alpha diversity metrics across yellow pea nodule microbiomes from spring (orange) and winter (blue) cultivars: Chao1 richness, Shannon diversity, Simpson’s diversity, and Pielou’s evenness. Center lines denote medians; box widths indicate interquartile ranges. No significant differences were observed between seasonal groups for any index (ANOVA and Kruskal–Wallis *p* > 0.05; n = 8 per group). **B)** NMDS ordination of microbial community composition. Non-metric multidimensional scaling (NMDS) based on Bray–Curtis dissimilarity shows the similarity of genus-level community profiles among individual samples. Samples are colored by seasonal cultivar group (orange = spring; blue = winter), with ellipses representing 95% confidence intervals. **C)** Principal Coordinates Analysis (PCoA) based on Bray-Curtis distance. PCoA illustrates variance in microbial composition across spring and winter samples. The first two principal coordinates explain 75.9% and 15.4% of variation, respectively. Dashed ellipses represent 95% confidence intervals for each group (n > 8 per seasonal cultivar). Differences in community composition were tested using PERMANOVA (Adonis2 function, vegan package) with 999 permutations. **D)** Alpha diversity comparisons by cultivar. Mean ± 95% confidence intervals of four alpha diversity metrics (Chao1 richness, Shannon diversity, Simpson diversity, and Pielou’s evenness) across individual cultivars. Each point represents the cultivar’s mean index value, colored by season (orange = spring; blue = winter). No statistical tests were applied at the cultivar level due to limited replicate counts (n = 3 per cultivar).

To explore cultivar-level patterns in more detail, we analyzed the alpha diversity indices (Shannon, Simpson, Pielou, Chao1) with 95% confidence intervals for each cultivar (**Figure 2D**). Although the confidence intervals for spring and winter cultivars broadly overlap, consistent with the non-significant Kruskal-Wallis tests, a few trends emerge. Winter cultivars (Keystone, Vail, MS-20W3) consistently show higher mean diversity and evenness in all four metrics. For example, MS-20W3 exhibits the highest Chao1 richness (mean approximately 60) and Keystone the highest Pielou’s evenness (mean approximately 0.71). The spring cultivars (Salamanca, Durwood, Carver) cluster at lower mean values (Chao1 approximately 40-55; Pielou approximately 0.48-0.58) (**Figure 2D**). These seasonal trends, although not statistically significant with the current replication (n = 2-3 per cultivar), suggest that winter varieties may harbor a broader and more balanced nodule microbiome.

Finally, we evaluated differential abundance at the genus level using both DESeq2 and Wilcoxon rank sum tests. After multiple testing corrections, no genera showed statistically significant differences between spring and winter samples (**Supplementary Table S3**). Although our hypothesis predicted seasonal cultivar-driven changes in the nodule microbiome, the lack of significant differences between diversity metrics and taxonomic profiles suggests a robust and conserved microbiome structure, largely dominated by *Rhizobium*, with any genotype-specific effects potentially masked by high variability between samples. In addition, the dominance of *Rhizobium* in all samples may have reduced the sensitivity of diversity metrics to detect differences between taxa of lower abundance. This effect likely contributes to the uniformity observed in the alpha and beta diversity indices and highlights the importance of complementary analyses that account for dominant taxa.

### Unmasking microbial diversity beyond *Rhizobium*

In all nodule samples, *Rhizobium* was consistently the most dominant bacterial genus, accounting for 84% to 98% of the total classified sequences. This dominance significantly compressed the diversity space, lowering the alpha and beta diversity estimates, which are highly sensitive to the evenness and distribution of taxa (Jousset et al., 2017; Willis, 2019). To address this, we filtered *Rhizobium* genus from the dataset and reanalyzed the microbial profiles. This filtering step enabled a more detailed examination of the less abundant, non-rhizobial taxa associated with nodules.

At the genus level, several taxa that were previously masked by *Rhizobium*’s dominance became more prominent, including *Crinalium*, *Tychonema*, *Rhodoligotrophos*, *Rickettsia*, and *Liberibacter* (**Figure 3A**). These genera were present in spring and winter cultivars and contributed more significantly to the overall structure of the community in the filtered dataset. Next, we recalculated species richness (Chao1), overall diversity (Shannon, Simpson), and community differences (Bray-Curtis) after *Rhizobium* removal. No significant effect of *Rhizobium* removal was observed on the total microbiome structure (*p* > 0.05; **Supplementary Figure S1**). The only difference was in evenness (Pielou index; *p* < 0.05; **Figure 3B**), with winter-type cultivars showing a more balanced read distribution between species.

**Figure 3.**
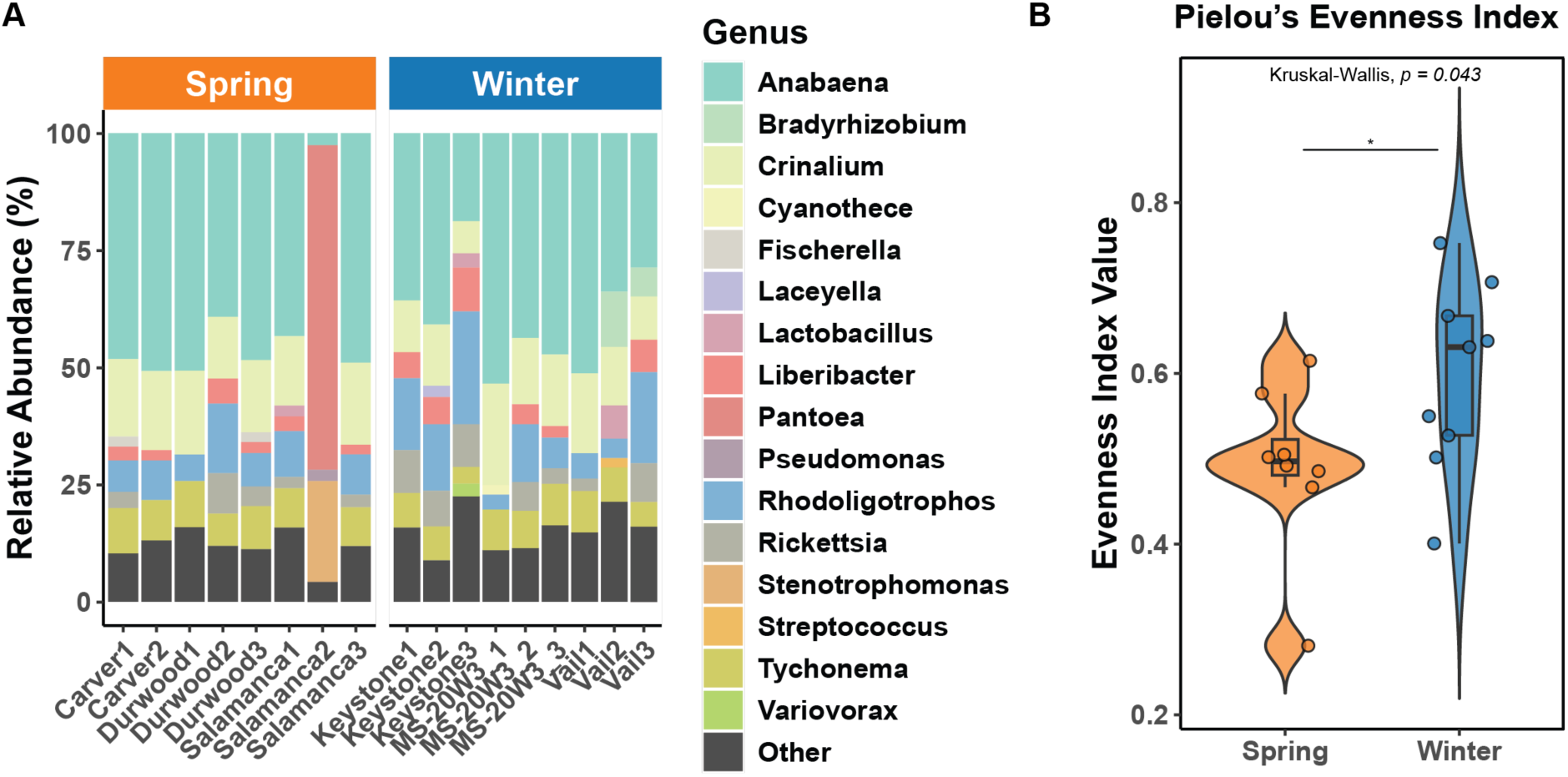
Revealing microbial community composition and evenness after *Rhizobium* sequence removal from yellow pea nodules. **A)** Genus-level composition of non-rhizobial nodule microbiomes. Stacked bar plots show the relative abundance (%) of bacterial genera in yellow pea nodules after computational removal of *Rhizobium* sequences. Samples are grouped by seasonal cultivar: spring (orange) and winter (blue). Remaining taxa span diverse genera, including *Anabaena*, *Crinalium*, *Liberibacter*, *Pantoea*, and *Rickettsia*, with notable shifts in relative contributions across cultivars. Bars represent individual biological replicates (n = 18 total), with each color corresponding to a specific genus as indicated in the legend. **B)** Increase in community evenness after *Rhizobium* removal. Violin plots display Pielou’s evenness index values for spring (n = 8) and winter (n = 9) samples following removal of *Rhizobium*. Evenness reflects how evenly taxa are distributed within each sample. Winter cultivars exhibited significantly greater evenness than spring cultivars (Kruskal–Wallis test, *p* = 0.043). Each dot represents one replicate; center lines indicate median values, and boxes reflect interquartile ranges.

### Core- and specific-microbiome in seasonal yellow pea cultivars

To explore microbial group-specific variation, we characterized the core and specific microbiome composition of yellow pea nodules from spring and winter cultivars at the species level after the removal of *Rhizobium* sequences. Core species were defined by absolute abundance-occurrence (Neu et al., 2021): taxa detected in at least 90% of cultivars within a seasonal group, with at least 500 reads in more than two replicates. We used raw read counts rather than relative abundance to minimize the impact of variable sequencing depths across samples, which can mask rare taxa and disproportionately emphasize dominant taxa. This classification revealed a structured microbial composition between winter and spring cultivars (**Figure 4A**; **Supplementary Table S4**). A total of 19 species were classified as part of the core microbiome shared between both seasonal groups, suggesting a conserved microbial baseline in yellow pea nodules. Although these 19 taxa represented only 9.7% of the total species richness, they accounted for 83% of total microbial abundance across samples (**Figure 4B**). Most of these taxa belonged to cyanobacterial lineages with known epibiotic lifestyles, including members of the Oscillatoriaceae, Nostocaceae, Hapalosiphonaceae, and Chroococcaceae families. Genera such as *Anabaena*, *Fischerella*, *Tychonema*, and *Gloeocapsa* are frequently associated with photosynthetic activity and have been observed in plant-associated environments (**Table 1**). The consistent presence of these taxa across cultivars suggests stable niche associations that can support or interact with nitrogen-fixing symbioses.

**Figure 4.**
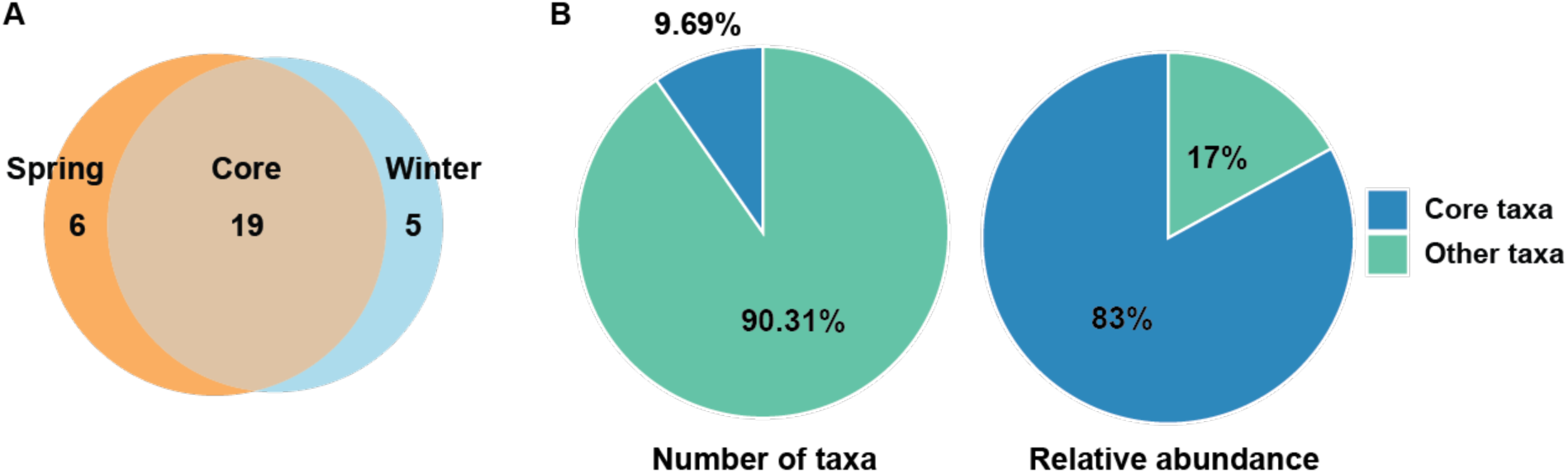
Core and group-specific non-rhizobial microbiome composition in seasonal yellow pea cultivars. **A)** Venn diagram summarizing the number of core taxa shared between spring and winter cultivars, as well as those exclusive to each seasonal group. **B)** Pie charts showing the proportion of core versus non-core (“other”) taxa by number of species (left) and total relative abundance (right). Although core taxa represent only 9.69% of species, they account for 83% of the total microbial abundance. *Notes:* Core taxa classification was based on prevalence (≥ 90% of samples in a group with ≥ 500 reads per taxon, with n > 2 replicates); relative abundances were averaged across all samples.

**Table 1.**
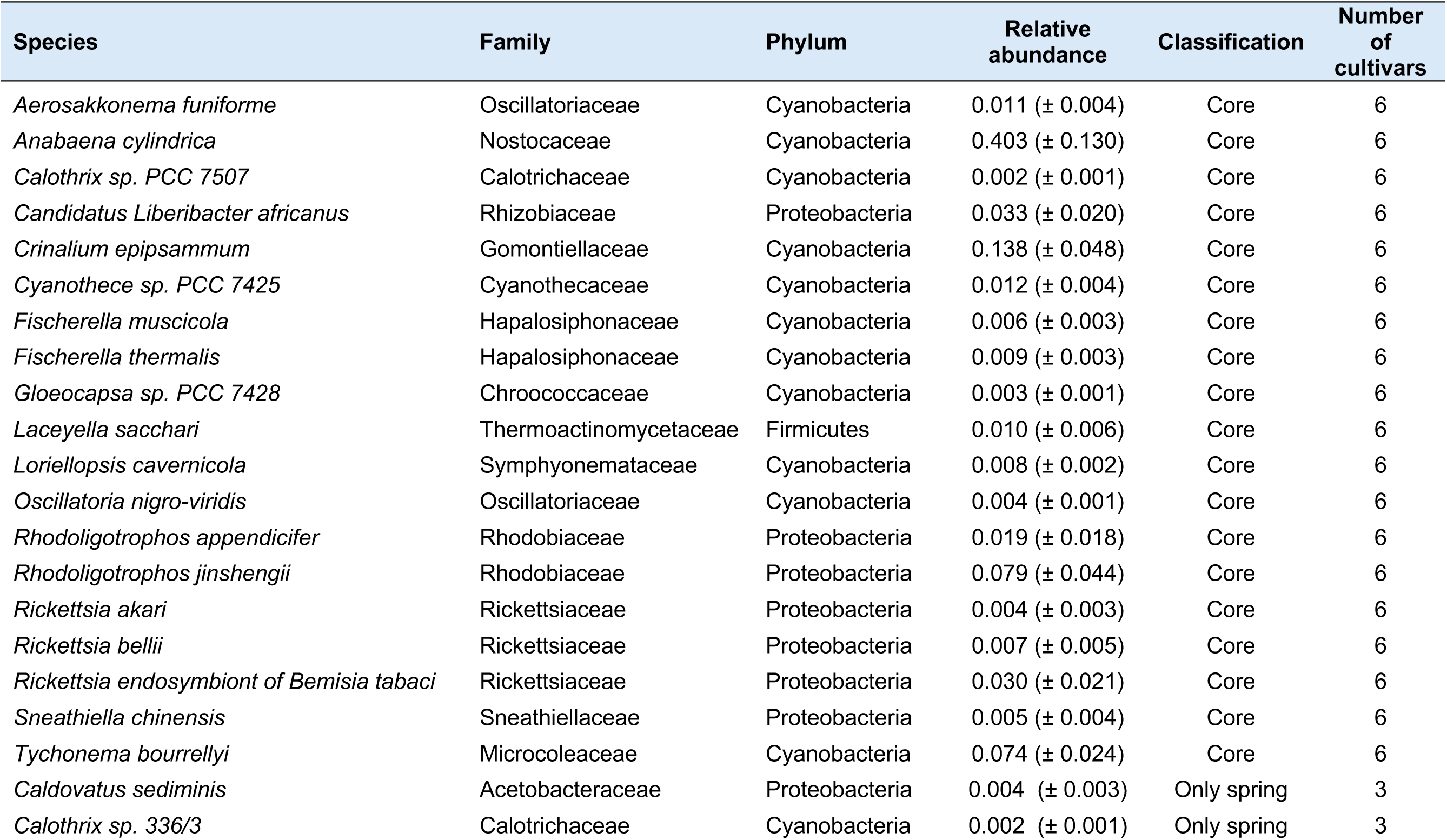
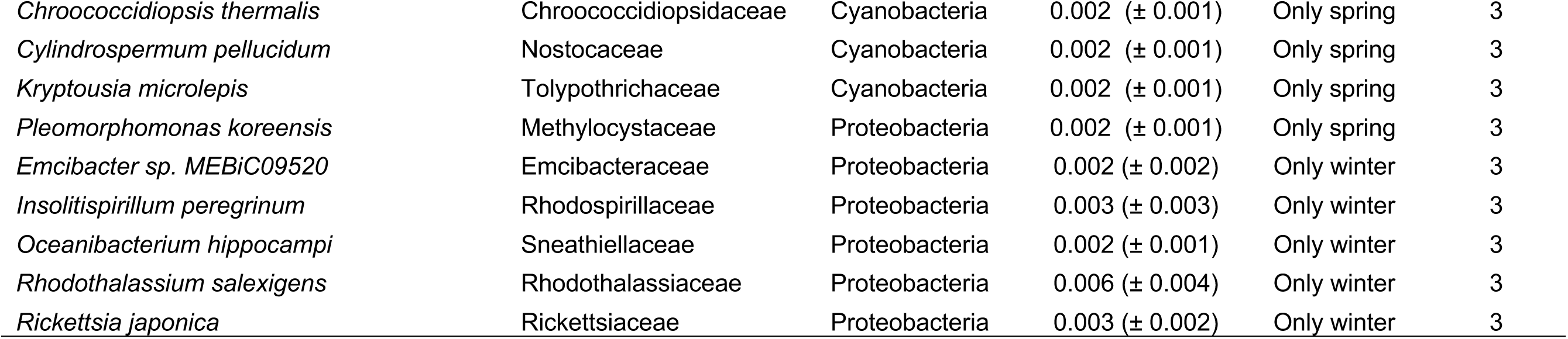
Taxonomic composition and relative abundance of core and seasonal-specific nodule-associated taxa.

To better understand seasonal specificity, we further examined the non-*Rhizobium* core taxa unique to each cultivar group. The spring-specific core microbiome included six species, mainly epibiotic cyanobacteria such as *Calothrix sp. 336/3*, *Cylindrospermum pellucidum*, and *Chroococcidiopsis thermalis*. These taxa are known for their resilience in variable environments and potential roles in nitrogen dynamics (Johansen et al., 2014; Baldanta et al., 2023; Uesaka et al., 2024). An additional spring-specific species, *Pleomorphomonas koreensis*, is a nitrogen-associated free-living bacterium previously isolated from plant rhizospheres, supporting its relevance in plant-microbe interactions (Im et al., 2016) (**Table 1**). The winter-specific core microbiome included five taxa, of which *Emcibacter sp.* (Meier et al., 2019), *Insolitispirillum peregrinum* (Balogavá et al., 2025), and *Rhodothalassium salexigens* (Madigan et al., 2025) have been identified in soil or aquatic environments and can play roles in carbon or sulfur cycling (**Table 1**). Although their specific functions in nodules remain unclear, their consistent presence suggests a possible adaptation to winter conditions and a contribution to the seasonal microbial structure. Together, these results reveal the presence of a dominant and stable non-*Rhizobium* core microbiome in yellow pea cultivars, accompanied by seasonal differences in cyanobacterial and free-living taxa.

### Functional insights from the core and group-specific microbiomes

To investigate the functional potential of microbial communities in yellow pea nodules, we applied FAPROTAX to species-level taxonomic profiles and grouped the predictions by microbial classification: (1) shared core microbiome, (2) spring-specific taxa, and (3) winter-specific taxa. The shared core microbiome exhibited a clear dominance of phototrophic functions, including general phototrophy, photoautotrophy, and oxygenic photoautotrophy, which showed the highest mean abundances among all predicted traits (**Figure 5B**; **Supplementary Table S5**). The functions associated with photosynthetic cyanobacteria and nitrogen fixation were also enriched in the core group, consistent with the presence of phototrophic and diazotrophic taxa, whereas traits such as intracellular parasitism were detected at lower abundance.

**Figure 5.**
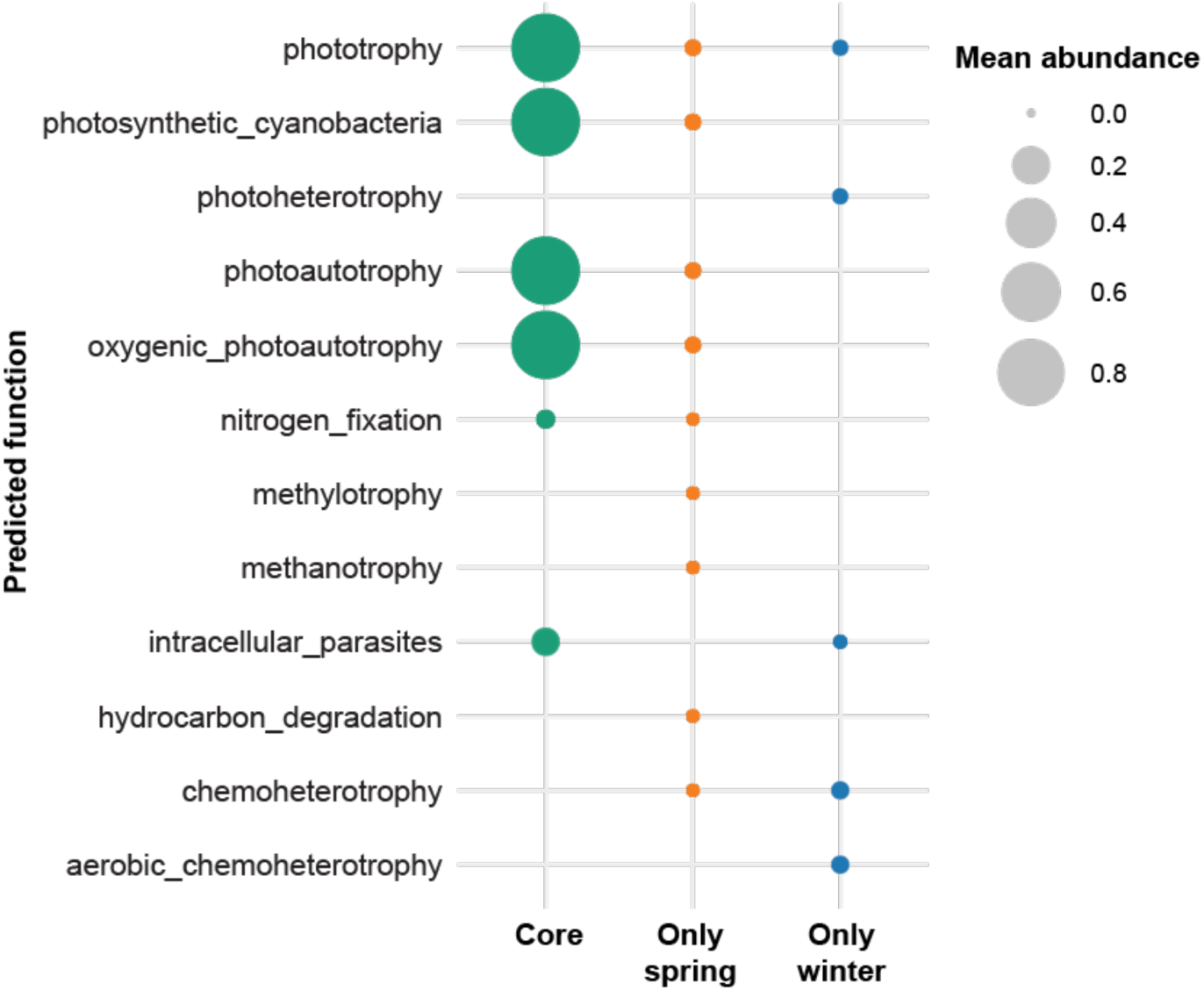
Predicted functional profiles of the nodule-associated microbiome across seasonal yellow pea cultivars. Bubble plot shows the mean relative abundance of predicted microbial functions for three groups of taxa: Core (detected in all cultivars), Only spring (exclusive to spring cultivars), and Only winter (exclusive to winter cultivars). Functional predictions were inferred using FAPROTAX based on taxonomic profiles. Bubble size reflects mean abundance per group, with larger bubbles indicating higher relative abundance.

In contrast, spring-specific taxa displayed a broader range of predicted functions, such as photoheterotrophy, methanotrophy, methylotrophy, and hydrocarbon degradation, but these occurred at lower relative abundance, suggesting a more functionally diverse but less dominant microbial contribution. Meanwhile, winter-specific taxa were primarily associated with aerobic chemoheterotrophy and photoheterotrophy, also at low abundance, indicating a more constrained metabolic repertoire. Altogether, these results highlight that the functional potential in yellow pea nodules is largely driven by a conserved core community, while season-specific groups contribute additional, but limited, functional variability.

## DISCUSSION

The advent of long-read sequencing technologies has substantially enhanced our ability to resolve complex microbial communities, particularly by improving taxonomic resolution in 16S rRNA-based studies (Tedersoo et al., 2021). However, each microbiome study presents unique characteristics that often require customized analytical workflows. This is especially relevant when considering the expected variability in microbial diversity between plant compartments. For example, the high microbial diversity typically found in the rhizosphere, phyllosphere, or floral tissues (Trivedi et al., 2020) contrasts with the lower diversity observed in specialized organs such as root nodules (Martinez-Hidalgo et al., 2017; Sharaf et al., 2019; Yurgel et al., 2025), which are often dominated by a single symbiotic taxon. In this study, we combined stringent filtering of host organellar reads and removal of *Rhizobium* sequences to disentangle the hidden diversity of the yellow pea nodule microbiome. This approach confirmed *Rhizobium leguminosarum* as the overwhelmingly dominant symbiont, while also exposing a structured assemblage of non-rhizobia taxa, including cyanobacteria and proteobacterial lineages, that form a reproducible core across cultivars.

### Dominant *Rhizobium* and implications for diversity patterns

We leveraged species- and genus-level resolution to enhance the detection of rare microbial taxa. As expected, the microbial profiles of the yellow pea nodules were overwhelmingly dominated by *Rhizobium*, consistent with its central role as the primary nitrogen-fixing symbiont in legumes (Mathesius, 2022). The high abundance of *Rhizobium* (84–98% of reads) reflects successful nodule colonization by compatible symbionts and is consistent with previous reports in peas and other legumes (Ji et al., 2017; Zhang et al., 2018; Yurgel et al., 2025). This dominance explains why richness (Chao1) remained stable between cultivars, whereas indices sensitive to abundance distribution (Simpson, Pielou) captured differences in evenness. In spring cultivars, higher evenness suggests that secondary taxa achieve more balanced representation, while winter cultivars appear more strongly structured by *Rhizobium* dominance. Similar findings in mung bean indicate that host genotype often influences not the absolute number of taxa but their relative balance, highlighting the selective filtering role of legumes (Favero et al., 2021). Such patterns underscore that diversity metrics must be interpreted considering symbiont dominance and the ecological specialization of nodules.

### Cultivar effects on nodule microbiome assemblage

Plant genetic variation exerts a significant influence on the assembly of microbiomes across different plant compartments, such as roots, nodules, and leaves (Singer et al., 2019; Brown et al., 2020; Ishida et al., 2022). Distinct genotypes create unique biochemical and physiological environments through differences in root exudation, tissue chemistry, or immune signaling, which in turn shape the recruitment, abundance, and persistence of specific microbial taxa (Zhang et al., 2023). This host-driven filtering effect helps explain why plants of the same species but with different genetic backgrounds often harbor contrasting microbial communities, even when grown under similar environmental conditions.

In our dataset, differences in *Rhizobium* abundance between cultivars were modest, but the consistent trend toward higher values in winter peas suggests genotype-associated recruitment. Parallel shifts also appeared in the non-rhizobial microbiome, plausibly reflecting subtle variation in exudate chemistry, immune tone, or nodule developmental timing (Zhalnina et al., 2018; Reid et al., 2018; Teixeira et al., 2021). Similar to reports in *Medicago truncatula*, our results reinforce that genotype effects are generally stronger in the rhizosphere and roots than in nodules (Brown et al., 2020). Soybean studies (Han et al., 2025) likewise demonstrate significant genotype influences in the rhizosphere, highlighting that the strength of this effect depends on plant compartment (Brown et al., 2020).

Although the magnitude of cultivar effects in our study was statistically modest, they point to a degree of host-driven filtering even within a single legume species, consistent with patterns observed across broader legume groups (Mutch et al., 2004; Hartman et al., 2017; Zhong et al., 2019). However, genotype-driven effects remain poorly documented in yellow pea, which crucially, nodules constitute a highly selective environment (Han et al., 2020), which likely compresses variation and makes cultivar effects more subtle.

### Non-rhizobial core: cyanobacteria and functional potential

We identified a conserved non-rhizobial core comprising primarily cyanobacteria (e.g., *Anabaena*, *Tychonema*, *Fischerella*), which are well documented in soil, biofilms, and plant-associated environments (Peng et al., 2019; Evseev et al., 2023). Their recurrent presence across cultivars suggests stable associations rather than sporadic colonization. Cyanobacteria possess traits such as oxygenic photosynthesis and diazotrophy that could complement nodule function, potentially buffering microaerobic conditions or supplying additional fixed nitrogen (Llamas et al., 2023). Whether they act as true endosymbionts or persist as epibionts on inner surfaces remains unresolved, but their consistent detection following rigorous sterilization indicates at least a close and reproducible association. Functional predictions reinforce this, with phototrophy and nitrogen fixation enriched in the shared core, while seasonal-specific taxa contributed additional but less abundant functions such as methylotrophy or sulfur cycling.

Similarly, the identification of non-rhizobial core and group-specific taxa with distinct predicted functional signatures, such as photoheterotrophy, methylotrophy, and hydrocarbon degradation, indicates that nodule-associated microbial communities may harbor functional diversity beyond canonical nitrogen fixation (see review by Bulgarelli et al., 2013). These observations parallel reports in peanut and soybean nodules, where non-rhizobial taxa extend the functional landscape beyond canonical symbiosis (Hossain et al., 2023; Han et al., 2020).

## CONSIDERATIONS AND IMPLICATIONS

While filtering strategies were effective in unmasking hidden diversity, limitations remain. Predictive functional profiling relies on taxonomic inference and cannot confirm the associated traits. Moreover, distinguishing between internalized bacteria and persistent epiphytes is challenging without microscopy or culture-based validation. In addition, the modest sample size constrains power to detect fine-scale genotype effects. Addressing these issues will require integrating amplicon sequencing with metatranscriptomics, imaging, and targeted inoculation assays to validate functional contributions of non-rhizobial taxa. Furthermore, the discovery of a stable, cyanobacteria-enriched core suggests that nodule microbiomes may provide ecological functions complementary to nitrogen fixation, with potential implications for crop resilience. Understanding whether these taxa enhance stress tolerance, modulate oxygen availability, or contribute to nutrient turnover could open avenues for microbiome-informed breeding or inoculant design. Future work should test whether core cyanobacteria confer measurable benefits to host performance and whether these associations are conserved across environments. Expanding comparative studies across legumes and seasonal ecotypes will clarify whether the observed patterns in yellow pea reflect a generalizable feature of legume symbioses.

## SUPPLEMENTARY DATA LIST

The following supplementary data are available in the online version of this article:

- **Supplementary Table S1:** NanoStat summary.
- **Supplementary Table S2:** Abundancy summary by taxon.
- **Supplementary Table S3:** DESEq summary analysis by genus (comparison between spring and winter groups).
- **Supplementary Table S4:** Non-rhizobial microbiome classification.
- **Supplementary Table S5:** Functional prediction (FAPROTAX).

## ACKNOWLEDGEMENTS

The authors extend their heartfelt gratitude to the dedicated team at Climate Smart Project and Graham Lab for their unwavering motivation. We also would like to express our deepest gratitude to the Research Cyberinfrastructure team at South Dakota State University for their invaluable support and resources. The high-performance computing infrastructure and technical assistance provided by the center were crucial for the completion of this research.

## AUTHOR CONTRIBUTIONS STATEMENT

HMD conceived and designed the bioinformatic approaches, analyzed the data, prepared figures and tables, authored or reviewed article drafts, and approved the final draft. JLGH supported the sequencing methodology, reviewed the analyses and article drafts, and approved the final draft. CG conceived the project and secured funding, discussed strategies and results, authored and reviewed article drafts, and approved the final draft.

## DATA AVAILABILITY

Full length 16S rRNA sequences were deposited in NCBI’s SRA under BioProject accession number PRJNA1310262.

## LIST OF FIGURES AND TABLES

**Figure S1.**
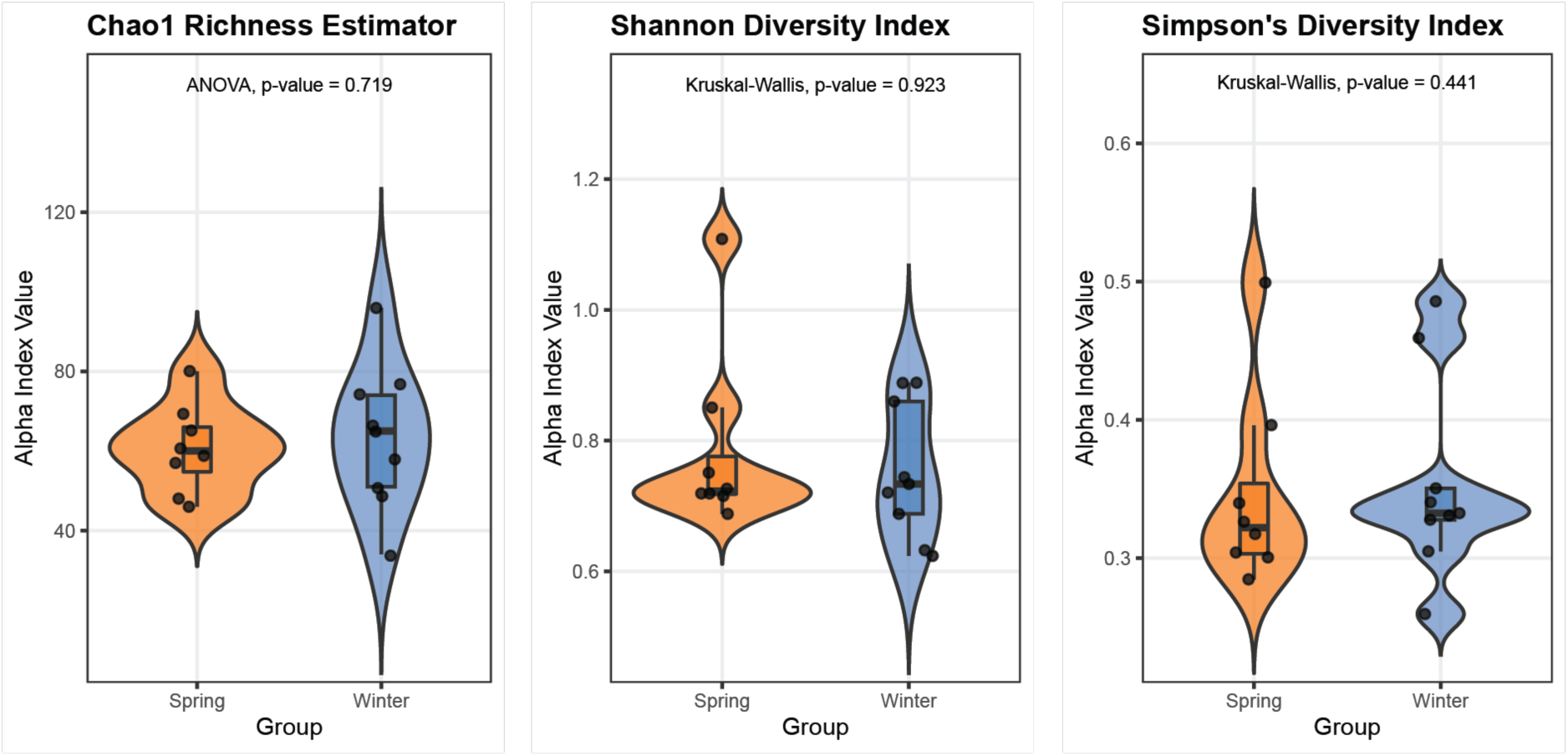
Seasonal cultivar effects on alpha and beta diversity of non-*Rhizobium* bacterial communities. Alpha diversity indices of spring and winter cultivars at genus-level. Violin plots show distributions of four alpha diversity metrics across yellow pea nodule microbiomes from spring (orange) and winter (blue) cultivars: Chao1 richness, Shannon diversity, and Simpson’s diversity. Center lines denote medians; box widths indicate interquartile ranges. No significant differences were observed between seasonal groups for any index (ANOVA and Kruskal–Wallis *p* > 0.05; n = 8 per group).

**Supplementary Table 1.** Summary of sequencing quality metrics generated by NanoStat before and after NanoFilt processing, including mean read length, mean read quality, number of reads, and N50 read length for each barcode.

**Supplementary Table 2.** Relative abundance of bacterial taxa summarized by phylum across all barcode-filtered samples, showing taxonomic profiles at higher classification levels.

**Supplementary Table 3.** Differential abundance analysis results generated with DESeq2, showing log_2_ fold changes, significance values, and taxonomic assignments for microbial taxa across cultivar groups.

**Supplementary table 4.** Core microbiome classification of bacterial species identified across samples, categorized as core, accessory, or specific to seasonal cultivar groups.

**Supplementary Table 5.** FAPROTAX-based functional predictions of microbial communities, showing the relative contribution of functional groups across all samples.

## LITERATURE CITED

Baldanta S, Arnal R, Blanco-Rivero A, Guevara G, Navarro Llorens JM (2023) First characterization of cultivable extremophile *Chroococcidiopsis* isolates from a solar panel. Frontiers in Microbiology 14:982422

Balogová, M., Šimonovičová, A., Vojtková, H., & Novák, O. (2025). The metabolic fingerprint of soil microbiota in an acidified environment using Biolog Microstationtm. Biologia, 1–15

Bever JD, Platt TG, Morton ER (2012) Microbial population and community dynamics on plant roots and their feedbacks on plant communities. Annu Rev Microbiol 66:265–283

Brown SP, Grillo MA, Podowski JC, Heath KD (2020) Soil origin and plant genotype structure distinct microbiome compartments in the model legume *Medicago truncatula*. Microbiome 8:139

Bulgarelli D, Schlaeppi K, Spaepen S, Van Themaat EVL, Schulze-Lefert P (2013) Structure and functions of the bacterial microbiota of plants. Annu Rev Plant Biol 64:807–838

Burghardt LT, Epstein B, Hoge M, Trujillo DI, Tiffin P (2022) Host-associated rhizobial fitness: dependence on nitrogen, density, community complexity, and legume genotype. Appl Environ Microbiol 88:e00526–22

Camargo AP, de Souza RS, Jose J, Gerhardt IR, Dante RA, Mukherjee S, Huntemann M, Kyrpides NC, Carazzolle MF, Arruda P (2023) Plant microbiomes harbor potential to promote nutrient turnover in impoverished substrates of a Brazilian biodiversity hotspot. ISME J 17:354–370

Carter KR, Nachtsheim AC, Dickman LT, Moore ER, Negi S, Heneghan JP, Sabella AJ, Steadman CR, Albright MB, Anderson-Cook CM et al. (2023) Drought conditioning of rhizosphere microbiome influences maize water use traits. Plant Soil 492:587–604

Crosbie DB, Mahmoudi M, Radl V, Brachmann A, Schloter M, Kemen E, Marín M (2022) Microbiome profiling reveals that Pseudomonas antagonises parasitic nodule colonisation of cheater rhizobia in Lotus. New Phytol 234:242–255

Curry KD, Wang Q, Nute MG, Tyshaieva A, Reeves E, Soriano S, Wu Q, Graeber E, Finzer P, Mendling W et al. (2022) Emu: species-level microbial community profiling of full-length 16S rRNA Oxford Nanopore sequencing data. Nat Methods 19:845–853

Daly C, Bryant K (2013) The PRISM climate and weather system—an introduction. PRISM Climate Group, Corvallis, OR

Dilla-Ermita CJ, Lewis RW, Sullivan TS, Hulbert SH (2021) Wheat genotype-specific recruitment of rhizosphere bacterial microbiota under controlled environments. Front Plant Sci 12:718264

Evseev P, Tikhonova I, Krasnopeev A, Sorokovikova E, Gladkikh A, Timoshkin O, Miroshnikov K, Belykh O (2023) Tychonema sp. BBK16 characterisation: lifestyle, phylogeny and related phages. Viruses 15:442

Favero VO, Carvalho RH, Motta VM, Leite ABC, Coelho MRR, Xavier GR, Rumjanek NG, Urquiaga S (2021) Bradyrhizobium as the Only Rhizobial Inhabitant of Mung Bean (Vigna radiata) Nodules in Tropical Soils: A Strategy Based on Microbiome for Improving Biological Nitrogen Fixation Using Bio-Products. Frontiers in Plant Science 11:602645

Galkiewicz JP, Kellogg CA (2008) Cross-kingdom amplification using bacteria-specific primers: complications for studies of coral microbial ecology. Appl Environ Microbiol 74:7828–7831

Hacquard S, Martin FM (2024) The chemical language of plant–microbe–microbe associations: an introduction to a virtual issue. New Phytol

Han Q, Ma Q, Chen Y, Tian B, Xu L, Bai Y, Chen W, Li X (2020) Variation in rhizosphere microbial communities and its association with the symbiotic efficiency of rhizobia in soybean. ISME J 14:1915–1928

Han X, Zhang X, Su M, Yang S, Feng S, Tian T, Chen N, Zhao G, Huang L, Wang G (2025) Genotype-specific rhizosphere microbiome assembly and its impact on soybean (Glycine max) yield traits. Front Sustain Food Syst 9:1518597

Hartman K, van der Heijden MGA, Roussely-Provent V, Walser J-C, Schlaeppi K (2017) Deciphering composition and function of the root microbiome of a legume plant. Microbiome 5:2

Hossain MS, DeLaune PB, Gentry TJ (2023) Microbiome analysis revealed distinct microbial communities occupying different sized nodules in field-grown peanut. Front Microbiol 14:1075575

Ishida JK, Bini AP, Creste S, Van Sluys MA (2022) Towards defining the core Saccharum microbiome: input from five genotypes. BMC Microbiol 22:193

Ji ZJ, Yan H, Cui QG, Wang ET, Chen WF, Chen WX (2017) Competition between rhizobia under different environmental conditions affects the nodulation of a legume. Syst Appl Microbiol 40:114–119

Johansen JR, Bohunická M, Lukesová A, Hrčková K (2014) Morphological and molecular characterization within 26 strains of the genus *Cylindrospermum* (Nostocaceae, Cyanobacteria), with descriptions of three new species. Journal of Phycology 50(1):187–202

Jousset A, Bienhold C, Chatzinotas A, Gallien L, Gobet A, Kurm V, Küsel K, Rillig MC, Rivett DW, Salles JF et al. (2017) Where less may be more: how the rare biosphere pulls ecosystems strings. ISME J 11:853–862

Liu F, Hewezi T, Lebeis SL, Pantalone V, Grewal PS, Staton ME et al. (2019) Soil indigenous microbiome and plant genotypes cooperatively modify soybean rhizosphere microbiome assembly. BMC Microbiology 19:201

Llamas A, Leon-Miranda E, Tejada-Jimenez M (2023) Microalgal and nitrogen-fixing bacterial consortia: from interaction to biotechnological potential. Plants 12:2476

Louca S, Parfrey LW, Doebeli M (2016) Decoupling function and taxonomy in the global ocean microbiome. Science 353:1272–1277

Love MI, Huber W, Anders S (2014) Moderated estimation of fold change and dispersion for RNA-seq data with DESeq2. Genome Biol 15:550

Madigan MT, Bender KS, Parenteau MN et al. (2025) Genomic highlights of the phylogenetically unique halophilic purple nonsulfur bacterium, Rhodothalassium salexigens. Extremophiles 29:12

Martínez-Hidalgo P, Hirsch AM (2017) The nodule microbiome: N2-fixing rhizobia do not live alone. Phytobiomes J 1:70–82

Mathesius U (2022) Are legumes different? Origins and consequences of evolving nitrogen fixing symbioses. J Plant Physiol 276:153765

Meier DV, Pjevac P, Bach W, Markert S, Schweder T, Jamieson J et al. (2019) Microbial metal-sulfide oxidation in inactive hydrothermal vent chimneys suggested by metagenomic and metaproteomic analyses. Environ Microbiol 21:682–701

Mutch LA, Young JPW (2004) Diversity and specificity of Rhizobium leguminosarum biovar viciae on wild and cultivated legumes. Mol Ecol 13:2435–2444

Neu AT, Allen EE, Roy K (2021) Defining and quantifying the core microbiome: challenges and prospects. Proc Natl Acad Sci USA 118:e2104429118

Oksanen J, Blanchet FG, Kindt R, Legendre P, Minchin PR, O’Hara RB, Simpson GL, Solymos P, Stevens MHH, Wagner H et al. (2013) Package ‘vegan’: community ecology package, version 2.5-7

Peng X, Bruns MA (2019) Development of a nitrogen-fixing cyanobacterial consortium for surface stabilization of agricultural soils. J Appl Phycol 31:1047–1056

R Core Team. (2021). R: A Language and Environment for Statistical Computing. Vienna, Austria: R Foundation for Statistical Computing. Retrieved from https://www.R-project.org/

Reid D, Liu H, Kelly S, Kawaharada Y, Mun T, Andersen SU, Desbrosses G, Stougaard J (2018) Dynamics of ethylene production in response to compatible Nod factor. Plant Physiol 176:1764–1772

Sergaki C, Lagunas B, Lidbury I, Gifford ML, Schäfer P (2018) Challenges and approaches in microbiome research: from fundamental to applied. Front Plant Sci 9:1205

Sharaf H, Rodrigues RR, Moon J, Zhang B, Mills K, Williams MA (2019) Unprecedented bacterial community richness in soybean nodules vary with cultivar and water status. Microbiome 7:63

Singer E, Bonnette J, Kenaley SC, Woyke T, Juenger TE (2019) Plant compartment and genetic variation drive microbiome composition in switchgrass roots. Environ Microbiol Rep 11:185–195

Sujkowska M, Górska-Czekaj M, Bederska M, Borucki W (2011) Vacuolar organization in the nodule parenchyma is important for the functioning of pea root nodules. Symbiosis 54:1–16

Tedersoo L, Albertsen M, Anslan S, Callahan B (2021) Perspectives and benefits of high-throughput long-read sequencing in microbial ecology. Appl Environ Microbiol 87:e00626–21

Teixeira PJ, Colaianni NR, Law TF, Conway JM, Gilbert S, Li H, Salas-González I, Panda D, Del Risco NM, Finkel OM et al. (2021) Specific modulation of the root immune system by a community of commensal bacteria. Proc Natl Acad Sci USA 118:e2100678118

Trivedi P, Leach JE, Tringe SG, Sa T, Singh BK (2020) Plant–microbiome interactions: from community assembly to plant health. Nat Rev Microbiol 18:607–621

Uesaka K, Fukuda N, Itoh K, Maruyama F, Hoshino T (2024) Restoration of the functional nif gene cluster by complex interspecies recombination in filamentous cyanobacteria. PLoS Computational Biology 20(3):e11249958

USDA (2022) Winter pea cover crop fact sheet. https://www.nass.usda.gov/sites/default/files/2024-07/Winter%20Pea%20Cover%20Crop%20Fact%20Sheet.pdf. Accessed 22 April 2025

USDA-GIPSA (2010) Official United States standards for grain: field peas. https://www.ams.usda.gov/sites/default/files/media/2010GIPSANR.pdf. Accessed 22 April 2025

Vann R, Reberg-Horton S, Castillo M, Mirsky S, McGee R (2018) Winter pea cultivar/breeding line screening for grain crop potential in the southeastern United States. Agron J 110:1217–1225

Vocanson A, Jeuffroy MH (2008) Agronomic performance of different pea cultivars under various sowing periods and contrasting soil structures. Agron J 100:748–759

Wendlandt CE, Regus JU, Gano-Cohen KA, Hollowell AC, Quides KW, Lyu JY, Adinata ES, Sachs JL (2019) Host investment into symbiosis varies among genotypes of the legume Acmispon strigosus, but host sanctions are uniform. New Phytol 221:446–458

Willis AD (2019) Rarefaction, alpha diversity, and statistics. Front Microbiol 10:2407

Wood DE, Lu J, Langmead B (2019) Improved metagenomic analysis with Kraken 2. Genome Biol 20:257

Yang HJ, Ye WW, Ze Y, Shen WL, Li SZ, Xing W, Chen JJ, Wang YC, Zheng XB (2023) Host niche, genotype, and field location shape the diversity and composition of the soybean microbiome. J Integr Agric 22:2412–2425

Yin C, Casa Vargas JM, Schlatter DC, Hagerty CH, Hulbert SH, Paulitz TC (2021) Rhizosphere community selection reveals bacteria associated with reduced root disease. Microbiome 9:86

Yurgel SN, McGee R (2025) Bacterial communities associated with food-quality winter pea cultivars grown in Pacific Northwest soils. Plant Soil

Zhalnina K, Louie KB, Hao Z, Mansoori N, Da Rocha UN, Shi S, Cho H, Karaoz U, Loqué D, Bowen BP et al. (2018) Dynamic root exudate chemistry and microbial substrate preferences drive patterns in rhizosphere microbial community assembly. Nat Microbiol 3:470–480

Zhang B, Du N, Li Y, Shi P, Wei G (2018) Distinct biogeographic patterns of rhizobia and non-rhizobial endophytes associated with soybean nodules across China. Sci Total Environ 643:569–578

Zhang R, Vivanco JM, Shen Q (2017) The unseen rhizosphere root–soil–microbe interactions for crop production. Curr Opin Microbiol 37:8–14

Zhong Y, Yang Y, Liu P, Xu R, Rensing C, Fu X, Liao H (2019) Genotype and rhizobium inoculation modulate the assembly of soybean rhizobacterial communities. Plant, Cell & Environment 42(6):2028–2044

Zhu YG, Peng J, Chen C, Xiong C, Li S, Ge A, Wang E, Liesack W (2023) Harnessing biological nitrogen fixation in plant leaves. Trends Plant Sci 28:1391–1405

